# Constructing well-defined neural networks of multiple cell types by picking and placing of neuronal spheroids using FluidFM

**DOI:** 10.1101/2024.09.03.610979

**Authors:** Sinéad Connolly, Katarina Vulić, Elaheh Zare-Eelanjegh, Marta Simonett, Jens Duru, Tobias Ruff, Blandine F. Clément, János Vörös

**Author notes:** Authors contributed equally (Sinéad Connolly, Katarina Vulić). Electronic supp. Information (ESI) available.

## Abstract

Controlled placement of single cells, spheroids and organoids is important for *in vitro* research, especially for bottom-up biology and for lab-on-a-chip and organ-on-a-chip applications. This study utilised FluidFM technology in order to automatically pick and place neuronal spheroids and single cells. Both single cells and spheroids of interest could be selected using light microscopy or fluorescent staining. A process flow was developed to automatically pick and pattern these neurons on flat surfaces, as well as to deposit them into polydimethylsiloxane microstructures on microelectrode arrays. It was shown that highly accurate and reproducible neuronal circuits can be built using the FluidFM automated workflow.

## 1 Introduction

With the emergence of human induced pluripotent stem cells lab-on-a-chip and organ-on-a-chip techniques are gaining greater importance in both biological and therapeutic research [1,2]. Constructing *in vitro* models of whole organs and systems on chips is becoming increasingly popular [3–6], and this trend is now expanding to encompass multi-organs on single chips [7,8]. Furthermore, individual units of biological systems can now be replicated *ex vivo* [9]. For example, simplified neuronal circuits of the nervous system have been successfully constructed, simulated, and analysed *in vitro* [10].

While these developments are exciting milestones toward individualised medicine and improved therapeutic testing techniques, they present specific challenges. Many require the precise positioning of cells, either to achieve a particular structure or to interact with their surroundings for stimulation or measurement purposes. For instance, in tumour cell migration experiments, metastatic cancer cells need to be positioned at a particular site away from the area of interest [11]. Similarly, in studies involving neuronal circuits, cells must be located as close to the electrodes of arrays as possible for successful stimulation and recording of electrical currents [12]. Consequently, there is a crucial need to accurately position spheroids, organoids, microtissues, and single cells within experimental microstructures to create well-defined, heterogeneous microphysiological systems that resemble physiological conditions.

This stipulation is particularly important for neuroscience. The brain remains one of the least-understood organs in the body, leading to a recent shift in focus from top-down neuroscience towards a bottom-up approach [10]. The aim of this approach is to achieve a more profound understanding of the brain’s functional units, which is expected to eventually provide insights into the entire organ. To support this goal, the growth of neurons for constructing neuronal circuits, the brain’s functional units, must be precisely controlled [13,14].

When neuron suspensions are deposited onto a surface, they tend to connect randomly, forming highly complex unidentifiable circuits that complicate the analysis of signal propagation. To address this issue, polydimethylsiloxane (PDMS) structures have been developed and mounted on multi-electrode arrays (MEAs). These structures guide axon growth, enabling the construction of reliable and reproducible circuits [12–19]. An example of such a microstructure is illustrated in Figures 1A and 1B) showing how cells can be seeded and used in such networks.

Although these techniques are well-established, they can still pose significant preparation challenges. Cells seeded in suspensions on top of PDMS microstructures do not always land in the designated wells. Instead, they may land on top of the microstructure, where they can grow and potentially compromise the integrity of the circuits beneath, particularly if they short-circuit them. Additionally, some wells may remain empty, reducing the number of viable experimental replicates and necessitating a larger sample size to achieve statistically significant results.

Moreover, when spheroids are manually seeded using a pipette, it is a manual process that needs to be conducted by the experimenter and it can be repetitive. While bioprinting [20] offers a potential solution to automate this process, current technologies struggle to print at the level of single spheroids, let alone single cells. This challenge is further complicated when working with flat surfaces at micron resolution or with microstructures, as ensuring the precise height of the placing surface is crucial to avoid damaging the placing device, the surface, or the biological samples. Especially, working with multiple cell types is cumbersome although it would be highly desirable for most organ-on-a-chip applications.

To address the aforementioned challenges, it was decided to incorporate fluidic force microscopy (FluidFM) into the preparation of neuronal circuits. This technique offers the precision needed to place cells accurately using gentle forces, overcoming the limitations of traditional methods and improving the reproducibility and efficiency of creating neuronal circuits.

FluidFM comprises an atomic force microscopy (AFM) cantilever, containing a hollow microfluidic channel [21– 43], see Figure 1C). Thus, the system merges the force-sensing abilities of AFM with the aspiration and dispensing abilities of a micropipette. This gives it a number of applications in material patterning and nanoprinting [44–69], injection into [70–74] and extraction from [73–78] cells and investigating mechanical properties such as stiffness [79–89] and adherence [42, 84, 86–123] of cells. Furthermore, because of its gentleness, accuracy and optical access, the FluidFM has also been used to pick and place cells [124–128]. The technique, outlined in Figure 1D), begins with the selection of the cell(s) of interest, from a cell-repellent substrate. The cantilever is lowered until contact is made with the selected spheroid or cell, detected using its AFM capabilities. A negative pressure is applied in the cantilever, and the object of interest is aspirated onto the cantilever’s aperture. This can then transfer the object to the location of interest, before depositing it there.

**Fig. 1:**
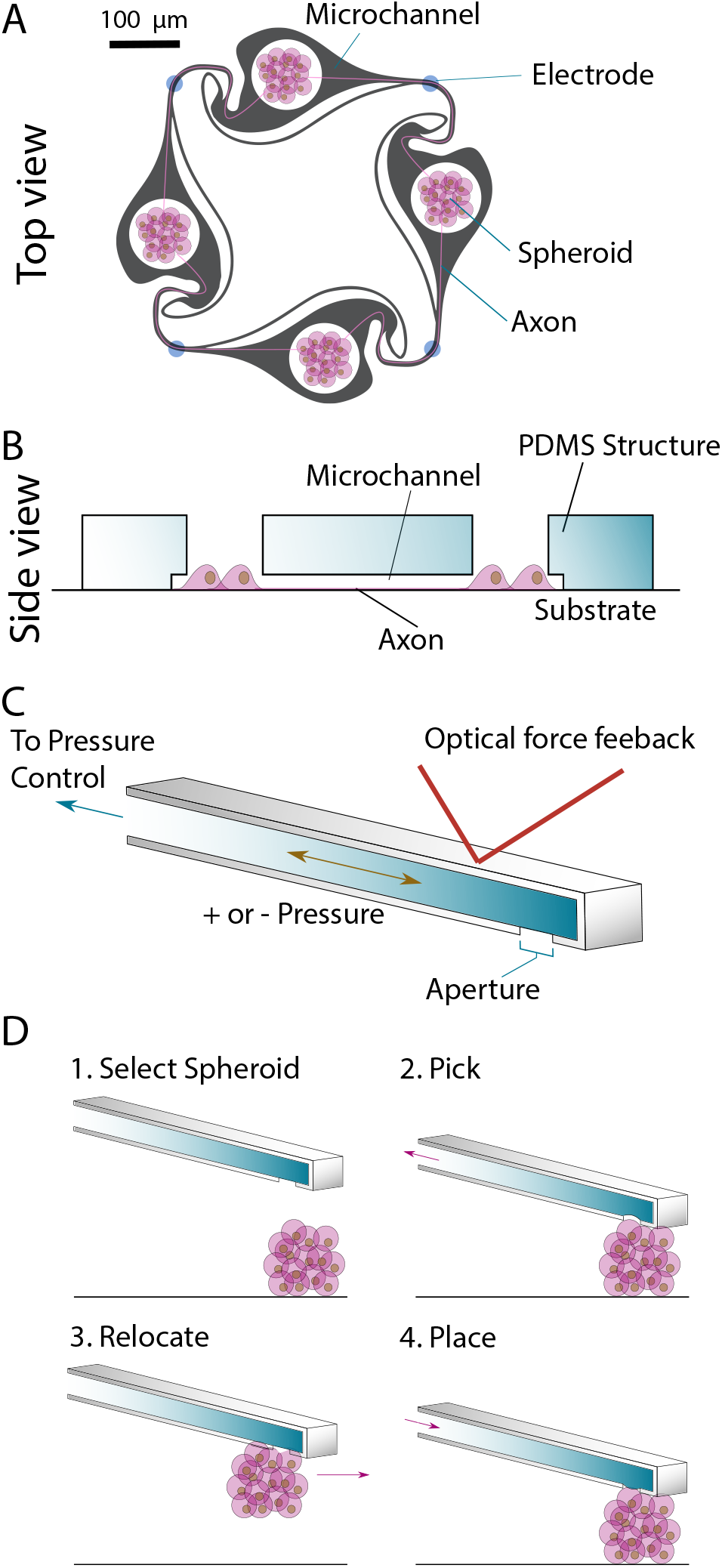
Picking and placing spheroids in microstructures. A) A top view of an example of a microstructure with spheroids seeded inside the wells. The shape of the microchannels encourages axons to grow in a clockwise direction only. [18] B) A side view of the microstructure containing cells. Wells are interconnected by axons. C) A fluidic force microscopy (FluidFM) cantilever. The hollow cantilever can be filled with a fluid in which a positive or negative pressure can be applied. D) Picking and placing spheroids using FluidFM. Negative pressure is applied through the cantilever to pick the spheroid from the surface and positive pressure is applied when placing the spheroid at the desired position.

Previously, the system has been used to pick and place bacteria [124,125], yeast [124,127] and mammalian cells including neurons [124, 127], HeLa cells [126, 128] and C2C12 cells [127]. Each of these studies focused on single cells, however, rather than on larger spheroids or organoids. Different studies investigated different parameters for picking and placing such as the picking substrate, probe coating and tip shape. Additionally, different characteristics such as size and level of adherence of different organisms required different parameters such as probe aperture size, aspiration and release pressures as well as the set-point forces. These parameters are outlined in Table 1.

**Table 1:**
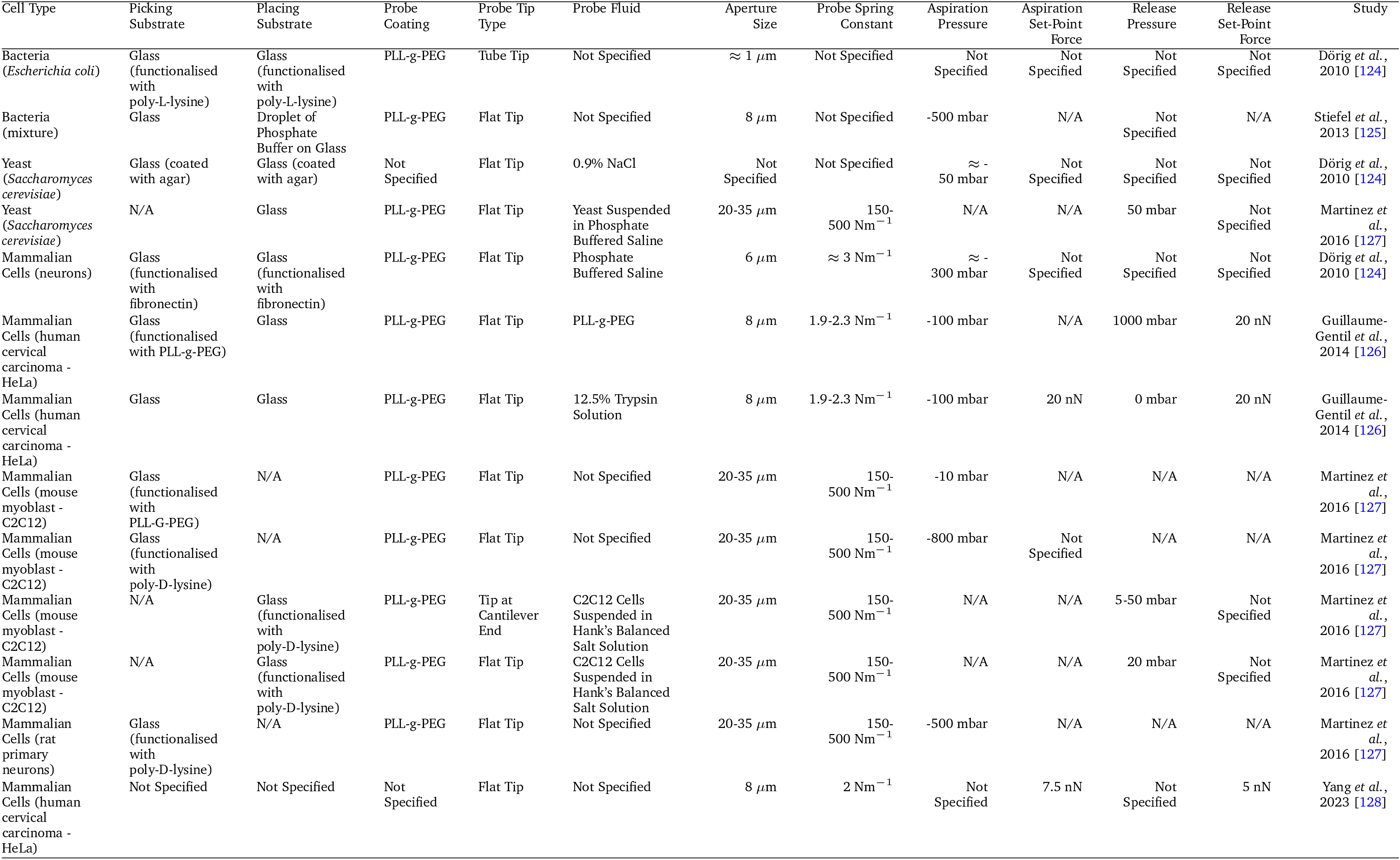
Previous parameters used in pick and place experiments

As previously noted, these studies utilised FluidFM for the precise picking and placing of single cells. However, until now, this technology has not been applied to multi-cellular organisms. Moreover, apart from the work by Yang *et al*. [128], the apparatus has been predominantly operated manually during the pick-and-place process, which is time-consuming and sub-optimal for preparing experimental structures.

The aim of this work is to employ FluidFM technology to construct functional neuronal networks, while automating the process to ensure that these networks can be built quickly and consistently. To achieve this, we used the FluidFM OMNIUM system to pick and place spheroids under various conditions, depositing them onto flat surfaces, MEAs and into the microstructures, and MEAs. Additionally, the process was largely automated using ARYA, the control software. This automation not only enables the automatic construction of neuronal microstructures using spheroids but also offers potential applications in other fields, as the methods described in this manuscript can be adapted for different cell types and microstructures.

## 2 Materials and Methods

### 2.1 Cell lines and spheroid formation

Two cell types; primary rat neurons and human induced pluripotent stem cell (hiPSC)-derived neurons were used in this study.

#### 2.1.1 Primary hippocampal neurons

Primary hippocampal cells were derived from E18 embryos of pregnant Sprague-Dawley rats (ETH Phenomics Center, Zürich, Switzerland). Cells were dissociated in house as previously described [18]. Briefly, embryonic brain tissue was placed in phosphate buffered saline (PBS), (10010-023, ThermoFisher, Zürich, Switzerland) with 22 *μ*M papain (P5306, Sigma-Aldrich & Cie, Buchs, Switzerland), 0.1% deoxyribonuclease (DNAse) (D5025, Sigma-Aldrich), 152 *μ*M bovine serum albumin (A7906, Sigma-Aldrich) and 10 mM D-(+)-Glucose (Y0001745, Sigma-Aldrich) in a water bath at 37 ^°^C for 15 minutes. The cells were then washed three times in NeuroBasal medium (NB) (21203-049, ThermoFisher) supplemented with 1% GlutaMAX (35050-061, ThermoFisher), 1% penicillin-streptomycin (15070-063, ThermoFisher) and 2% B-27 supplement (17504-044, ThermoFisher), which is denoted as complete NB. The first wash was additionally supplemented with 10% fetal bovine serum (FBS) to neutralize the action of the papain. Following dissociation, cells were maintained in complete NB.

#### 2.1.2 hiPSC-derived neurons

Neurogenin2 (NGN2) expressing hiPSC-derived neurons (Novartis, Basel, Switzerland) were obtained in cryogenised aliquots and stored in liquid nitrogen until use. Cells were thawed as previously described [14]. Briefly, a cryopreserved aliquot was removed from the liquid nitrogen and placed in a water bath at 37 ^°^C to thaw. The thawed cell solution was then added slowly to 4 mL of warmed neurobasal medium supplemented with 1% GlutaMAX, 1% penicillin-streptomycin, 2% B-27 supplement, 1% N-2 supplement (17502048, ThermoFisher), 0.1% BDNF recombinant protein (450-02-1MG, ThermoFisher) and 0.1% GDNF recombinant protein (450-10-1MG, ThermoFisher), the solution was centrifuged and the supernatant was replaced with NB differentiation medium (NBD). Following thawing, cells were maintained in NBD.

Following both cell dissociation and cell thawing, cells were counted and their viability was evaluated using a cell counter (Countess, ThermoFisher). Both cell types were cultured in an incubator at 37 ^°^C and 5% CO_2_.

#### 2.1.3 Spheroid formation

Spheroids were formed using AggreWell 400 microwell culture plates (34415, STEMCELL Technologies, Grenoble, France). The spheroid-forming wells were prepared prior to cell seeding by adding 0.5 mL of anti-adherence rinsing solution (07010, STEMCELL Technologies) to each well, and then centrifuging the plate to remove any bubbles from the microwells. The anti-adherence rinsing solution was then removed from the well and fresh, warmed medium was added. Cells were used immediately following either dissociation or thawing, and were added to wells at predetermined concentration, depending on the desired number of cells per spheroid or the desired spheroid size. The AggreWell plate was centrifuged again at a lower speed, and cells were left in the incubator for at least one night overnight to form spheroids. Spheroids were always used within one week of formation.

### 2.2 Substrate preparation

#### 2.2.1 PDMS microstructures

PDMS microstructures were designed and prepared as described previously [12–14]. Briefly, microstructures were drawn in Python, and then fabricated on a wafer using a standard soft lithography process [18] (Wunderlichips GmbH, Zürich, Switzerland). An example of such a microstructure is shown in Figures1 A) and B), however, as a proof of concept, different designs with different parameters were used in these experiments. Prior to mounting the PDMS structures on glass dishes (see section2.2.2) and MEAs (see section2.2.3), the desired microstructure was cut from the wafer, and placed bottom up on a glass slide. The base of the microstructure was then plasma cleaned (PDC-32G (18 W), Harrick Plasma, New York, USA) prior to mounting.

#### 2.2.2 Glass-bottom dishes

Round glass slides (10343435, ThermoFisher) were mounted onto WillCO dishes (KT-3522, WillCO Wells, Amsterdam, The Netherlands) according to the manufacturer’s instructions. Dishes were ultrasonicated, with isopropanol, and cleaned with isopropanol and ultrapure water (Milli-Q, Sigma-Aldrich). The dishes were blow dried using nitrogen and plasma cleaned. For experiments involving placing cells on flat surfaces, dishes were then coated with a solution of 0.1 mgmL^−1^ poly-D-lysine (PDL) (P6407, Sigma-Aldrich) in PBS for 45 minutes. The solution was then removed and the dish was rinsed three times with ultrapure water. The dish was then filled with fresh media and kept in the incubator until ready for use.

For experiments involving placing cells in microstructures, glass dishes were prepared as described above, up to and including the plasma cleaning step. Following this, the microstructure was placed top up on the glass base of the dish. In order to aid in placing the microstructure as flat as possible, a drop of isopropanol was placed on the microstructure, allowing the structure to be manipulated easily without any friction between it and the glass slide which would have caused it to tear. The dish was then left to sit for at least an hour to allow the microstructure to adhere properly to the glass dish. Next, the dish was filled with PDL and placed in a desiccator for an hour in order to remove any air bubbles that were trapped in the channels or the wells of the microstructure. The solution was then removed and the dish was rinsed three times with ultrapure water. The dish was then filled with fresh media and kept in the incubator until ready for use.

#### 2.2.3 Microelectrode arrays

Standard microelectrode arrays (60MEA500/30iR-Ti-gr, Multi Channel Systems, Reutlingen, Germany) were prepared as placing substrates. MEAs were plasma cleaned along with PDMS microstructures as outlined above. Following this, the microstructure was placed top up on the base of the MEA. In order to aid in placing the microstructure as flat as possible, a drop of isopropanol was placed on the microstructure, allowing the structure to be manipulated easily without any friction between it and the glass slide which would have caused it to tear. Additionally, this allowed the microstructure to be aligned with the electrodes of the MEA for electrical recordings of axons. The MEA was then left to sit for at least an hour to allow the microstructure to adhere properly to the MEA. Next, the MEA was filled with PDL and placed in a desiccator for an hour in order to remove any air bubbles that were trapped in the channels or the wells of the microstructure. The solution was then removed and the MEA was rinsed three times with ultrapure water. The MEA was then filled with fresh media and kept in the incubator until ready for use.

### 2.3 Experimental apparatus

#### 2.3.1 FluidFM OMNIUM

Pick and place experiments were conducted using the FluidFM OMNIUM, which consists of an AFM connected to a pressure-controlled microfluidics system (Cytosurge AG, Zürich, Switzerland). The system is surrounded by an incubator, with a circulating airflow filtered by a high-efficiency particulate air (HEPA) filter, and all experiments were conducted at 37 ^°^C and 5% CO_2_. The system is operated using the control software, ARYA (Cytosurge AG), which interfaces all the components of the system including the pressure controller, pump, microscope, control head and stage. This software also has a ‘macros’ function that allows actions to be pre-programmed to allow for automated experiments.

#### 2.3.2 Probes and probe preparation

Two types of probes were used; flat tips and tube tips. The flat tipped probes (CYPR/001558, Cytosurge AG) are made of silicon nitride and have an 8 *μ*m aperture diameter and 2 Nm^−1^ spring constant, see Figure 4A left. They are commercially available, and were purchased ready for use. The aperture is located on the underside of the probe and the rim of the aperture is flush with the base of the probe. The tube tipped probes were purchased externally from (SmartTip BV, Enschede, The Netherlands), they were inspected in house before being glued to the probe holder by Cytosurge AG. Tube tipped probes had an aperture diameter of 10-14 *μ*m and a spring constant of 4 Nm^−1^. The tube tipped probes were shaped in much the same way as the flat probes, however they had an additional protrusion (or tube) extending down from the aperture on the cantilever of 20 *μ*m, see Figure 4 A) right. Prior to experimentation, probes were plasma cleaned, and in some experiments coated to prevent cell adherence to the probe. Probes were either coated with PAcrAm (PAcrAm-g-(PMOXA,NH2,Si), SuSoS, Zürich, Switzerland) or with Sigmacote (SL2, Sigma-Aldrich). Probes coated with PAcrAm were filled with 1 *μ*L of the desired liquid (either water-based solution or oil), immersed in PAcrAm diluted in ultrapure water, for at least 30 mins, before being rinsed three times in PBS. Probes coated with Sigmacote required more steps. Probes were placed in a specialised container with 25 *μ*L of Sigmacote solution, which did not come into direct contact with the probe. The container was subsequently closed and placed inside a desiccator. The container was designed such that when a negative pressure was applied, the Sigmacote vapour was pulled through the probe, therefore coating the inside, as well as the outside of the probe. The probe was coated overnight, and placed in the oven at 60 ^°^C for 1 h the next day. Flat tipped probes were considered to be single use, and were discarded after each experiment. Tube tipped probes were reused, and stored in PBS at 4 ^°^C between experiments. Reused probes were refilled prior to experimentation.

#### 2.3.3 Experimental setup

Prior to experimentation, cells and placing substrates were prepared as outlined in Sections 2.1 and 2.2 above. Following mounting of the microstructures, the placing locations or the placing wells were programmed to be used in ARYA using the FluidFM Plate Editor (Cytosurge AG). Points were placed in a specific formation according to the design of the PDMS structure in either the glass-bottomed dish or the MEA using the FluidFM Plate Editor, and could later be aligned with the actual dish in ARYA to correct for distortions in the microstructure, or for a different alignment of the microstructure in the glass-bottomed dish or MEA.

The probe coating was carried out as described above in Section 2.3.2, and the probe was set up using the ‘Preparation Advanced’ function in ARYA.

Finally, the macros steps were set up. These parameters may have differed slightly, depending on the investigation, but in short, a number of picking sites and placing sites were manually selected using the ARYA software. Once the macros were set-up, the cantilever approached the first spheroid until a setpoint of 300 nN was reached. A negative pressure of -400 mbar was applied in the probe for 2 s. The cantilever was raised and moved horizontally to the placing site, where it was lowered until it reached a setpoint of 600 nN. Here, a positive pressure of 1000 mbar was applied for 20 s. The cantilever was then raised and moved to the picking location of the second spheroid, where the cycle began again until all picking and placing was completed. Every three cycles, the probe was ‘washed’ using 4% Tergazyme (Z273287, Sigma-Aldrich) solution, These steps are described in greater detail in the supplementary information S1 and S2 for spheroids and single cells respectively.

## 3 Results and Discussion

We show that FluidFM can be used to successfully pick and place (P&P) neural spheroids and single neurons onto various surfaces, including glass coated with cell-adhesive materials and into PDMS wells adhered to microelectrode arrays or glass-bottom dishes. Since FluidFM OMNIUM is equipped with an incubator, multiple hours P&P experiments were possible if needed. The experiments demonstrated successful P&P for rat primary neurons and hiPSC-derived neurons. For spheroids, the best results were achieved two days after spheroid preparation, whereas for single cells, optimal conditions were established immediately upon thawing or dissociating the cells. Multiple experimental conditions were optimised for P&P, which will be detailed in the following subsections.

### 3.1 Tuning probe parameters to facilitate spheroid placement

One of the main challenges in P&P is preventing unwanted adhesion of picked cells to the probe cantilever, which can impede their successful placement. This is of particular importance with neurons which tend to be ‘stickier’ than other cell types. For effective picking, a spheroid or single cell must adhere to the probe cantilever just enough to be transferred from the picking dish to the placing dish. However, the adhesion must be weak enough that the cell or spheroid can detach from the cantilever when positive pressure is applied and the cell-adhesive substrate is engaged, allowing the cell or spheroid to remain on the placing dish. Several probe parameters can be optimised to enhance cell placement and increase the placement rate. These parameters include probe filling, probe coating, and probe shape, each of which will be discussed in the following sections.

#### 3.1.1 Successful P&P obtained both with water-based and oil probe filling

Neural spheroids could be P&P successfully using two types of probe fillings: water-based and oil. Water-based filling involved using 1 *μ*L of ultrapure water, PBS, or NB. It was noticed that partial attachment of a cell to the cantilever allowed the water-based solutions to flow around it. Therefore, we investigated whether an oil filling would improve cell detachment due to the presence of a stable oil-water interface. Figure 2 A) showcases examples of successful P&P for spheroids of various sizes on different placing substrates with both water-based and oil fillings. In these experiments, the external probe surface (that is in direct contact with the spheroid or cell) was not coated.

**Fig. 2:**
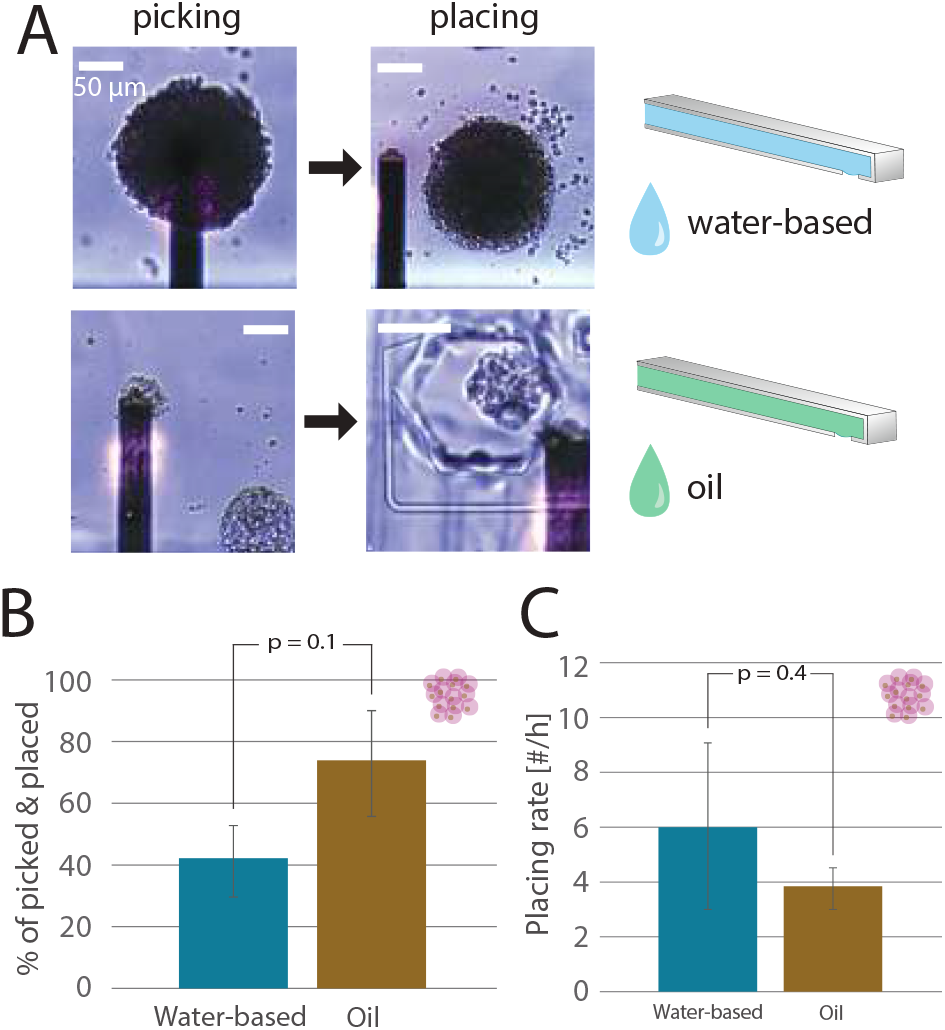
P&P of spheroids with different probe fillings. A) Demonstration of successful P&P for smaller and larger spheroids onto a glass surface and inside the PDMS well. B) Percentage of picked spheroids successfully placed on the substrate by differently filled probes. C) Rate of spheroid placement for differently filled probes.

In Figure 2 B), oil probe filling appears to offer a more successful placement for picked spheroids, with a mean value slightly above 40% for water-based filling compared to roughly 75% for oil filling. However, statistical analysis shows no significant difference in the percentage of spheroids placed on a substrate between the two probe fillings. Conversely, Figure 2 D) shows a higher mean placing rate for water-based filled probes, but also a higher standard error of mean, resulting again in no significant difference between the two filling types. We observed during experimentation, that probes with an oil filling tended to block easily and required repeated pressure changes and washing steps to clear such blockages, difficulties which did not tend to happen in the water-based fillings. This may explain the apparent discrepancy between the two. Despite this, the results suggest that the oil-filled probe was a more reproducible choice.

Therefore, we concluded that probe filling is not a crucial parameter for reliable and fast placement of picked spheroids. Consequently, in subsequent experiments, we used either water-based or oil-based fillings, varying other parameters. Notably, for single cells, no successful placements were achieved using oil-filled probes, indicating that water-based filling is the better choice for single-cell placement.

#### 3.1.2 Higher placing rate obtained with PAcrAm than with Sigmacote as probe coating

The next parameter to consider in achieving more reproducible spheroid placement on a substrate was coating the external surface of the probe. We believed that making the area that is in contact with the picked spheroid anti-adhesive in combination with a cell-adhesive substrate would facilitate spheroid detachment from the probe, while the negative pressure applied inside the cantilever would be enough to keep the spheroid attached upon transfer. Additionally, it had been noted in the previous set of experiments, that oil tended to form pockets inside the probe, leading to probe blockages, and extended cleaning times. We therefore anticipated that coating the probes containing an oil filling with a hydrophobic coating would improve the movement of oil within the probe and reduce blockages, therefore improving the placement rate of the spheroids. The anti-adhesive coatings tested were PAcrAm and Sigmacote. PAcrAm is a hydrophillic, surface-adhesive polymer, resistant to cell adhesion. Glass substrates are coated within 30 min with a monolayer by dipping the substrate in an aqueous solution, and rinsing afterwards [129]. Sigmacote is a solution of a chlorinated organopolysiloxane in heptane that readily forms a covalent, hydrophobic, microscopically-thin film on glass. Similarly to PAcrAm, it has been used as a protein- and cell-repellent coating. [130] Sigmacote is applied as a vapour solution. In the experiments where probe coatings were compared, we filled the probe with oil.

In Fig. 3 A), we report a significantly higher percentage of picked spheroids that were placed at the target spot when the probe was coated with PAcrAm. The percentage is slightly below 70%, which is comparable to the percentage when using a non-coated probe filled with oil (shown in Fig.2 B)). Similarly, the performance of a PAcrAm-coated probe exceeds the performance of a Sigmacote-coated probe in terms of placement rate, which can be observed in Figure 3 B). The placement rate with a PAcrAm-coated probe is again comparable with an oil-filled non-coated probe, indicating that using a coated probe does not lower the performance, but also does not significantly improve cell detachment during placement. We attribute the over-performance of PAcrAm-coated over Sigmacote-coated probes to the fact that PAcrAm forms a a few nm thick monolayer at the probe surface as opposed to a micrometer Sigmacote layer. Thicker layers might clog the aperture of the probe, which in the case of the flat probe described in 2.3.2, corresponds to 8 *μ*m, thus hindering the liquid flow through the cantilever. Additionally, it was noted during experimentation that the combination of oil with a Sigmacote-coated probe was so anti-adhesive, that it repelled spheroids away from the cantilever on approach, regardless of the negative pressure applied. This also explains the exceptionally low placing rate. Both the P&P percentage and the placement rate in the case of PAcrAm-coated probes are comparable with the non-coated probe filled with oil, not affecting the oil flow. While Sigmacote might improve the flow of oil in the probe, potential clogging of the probe presents a difficulty.

**Fig. 3:**
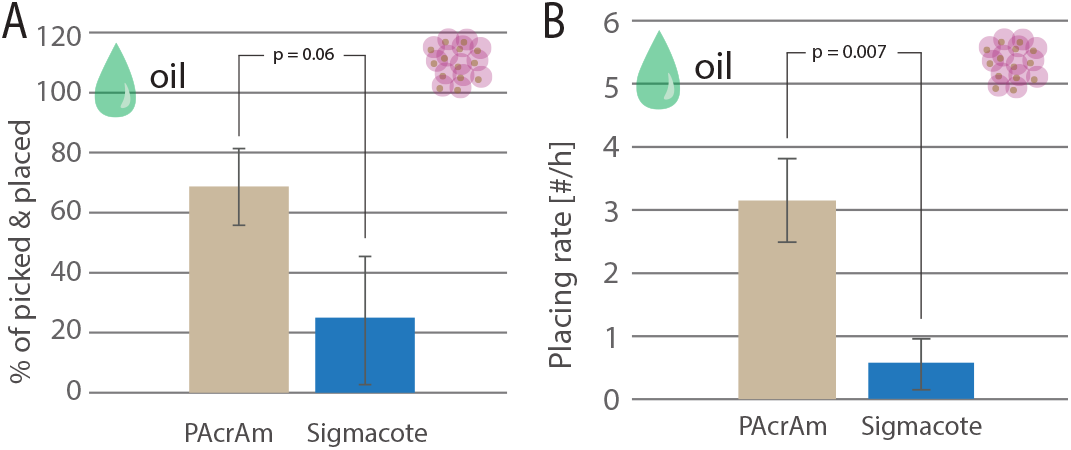
P&P of spheroids with different probe coatings. A) Percentage of picked spheroids successfully placed on the substrate by differently coated probes. B) Rate of spheroid placement for differently coated probes.

**Fig. 4:**
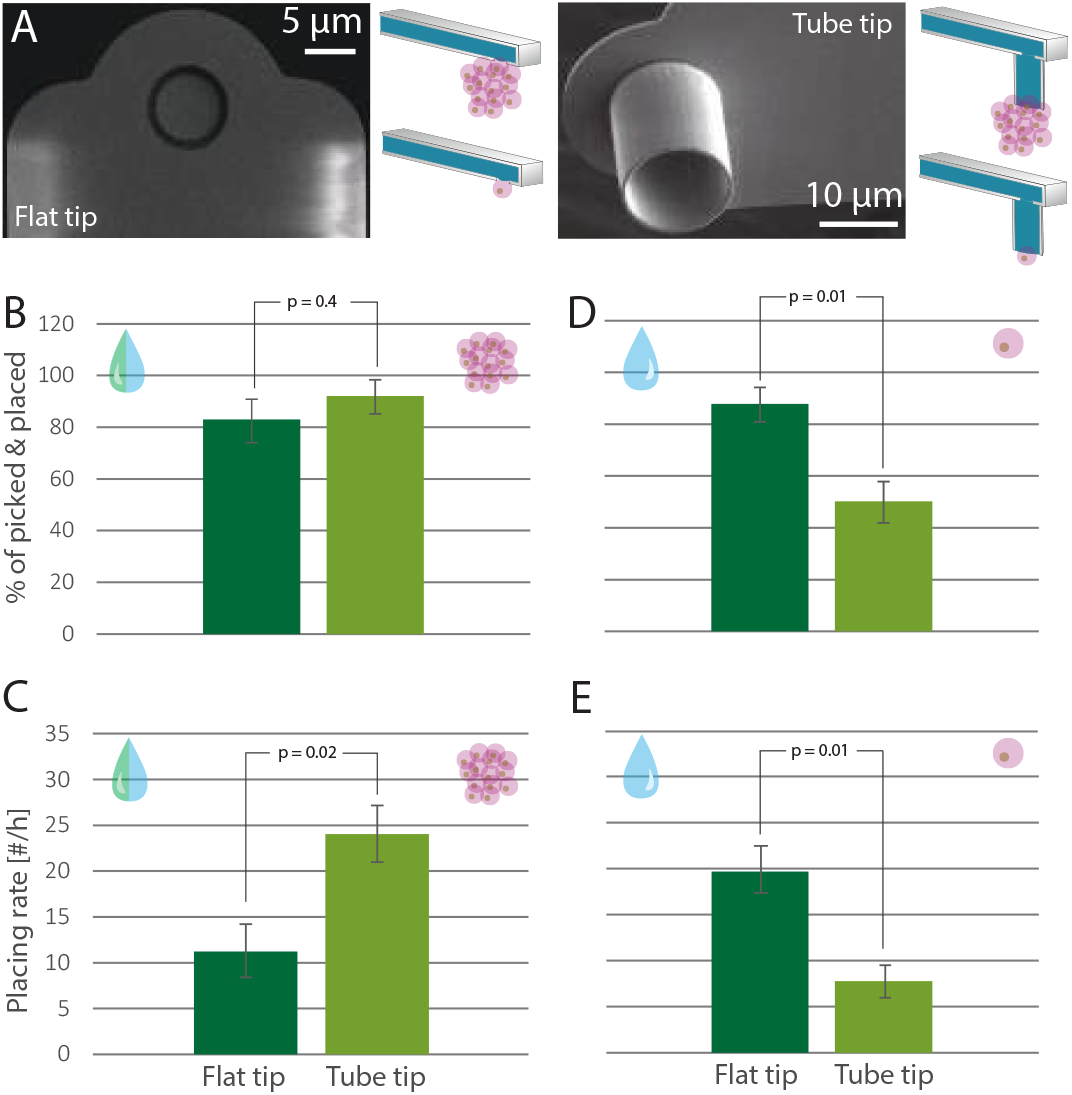
P&P of spheroids and single cells with differently-shaped probes. A) The geometry of the flat-tipped (left) and tube-tipped (right) probe. B) Percentage of picked spheroids successfully placed on the substrate by differently shaped probes. C) Rate of spheroid placement by differently shaped probes. D) Percentage of picked single cells successfully placed on the substrate by differently shaped probes. E) Rate of single cell placement by differently shaped probes.

#### 3.1.3 Tube-tipped probe significantly improves spheroid placement rate

The last probe parameter we tested to improve the P&P percentage and the placement rate of neural spheroids and single neurons was the probe shape. We compared picking and placing of spheroids and single cells using flat-tipped and tube-tipped probes. Since unwanted attachment of the cells to the probe is the main reason for unsuccessful placement, this was specifically targeted throughout this study. Though it was anticipated that the use of oil in the probe would aid in this respect, it created other problems with probe blockages. Therefore, an alternative approach was considered of reducing the contact area between the probe and the cell or spheroid, which was expected to improve the situation. While the shallow flat tip likely makes contact with the cells throughout the area of the opening, the tube tip only contacts the cells along its perimeter. In addition, the tube tip also prevents direct cell attachment to the cantilever around the opening. The geometry of the probes is described in detail in Section 2.3.2 and shown in Figure 4 A).

So far, the best method for picking and placing spheroids was deemed to be the use of an oil filling in the probe without the need for a coating. However, as the tube-tip would have a reduced contact area with the spheroid, it was believed, therefore, that oil would not be necessary as a filling in the tube-tips. Therefore, a comparison is made here between flat tips with an oil filling and tube-tips with a water-based filling. In Figure 4 B), we observed that the success rate of placing picked spheroids is similarly high for both oil-containing flat-tipped and PBS-containing tube-tipped probes, with mean percentages exceeding 80% in both cases. However, the significant improvement lies in the rate at which spheroids are picked and placed at the target location, as shown in Figure 4 C). Here, the tube-tipped probes demonstrate a much higher placement rate, more than double that of the flat-tipped probe. This is likely due to the reduced time spent cleaning the probe or attempting to remove oil pockets from inside the probe. The mean placement rate approaches 25 spheroids per hour, indicating that probe shape has the most substantial impact on placement rate among the three probe parameters considered: probe filling, probe coating, and probe shape. Unlike the other two parameters, which involve surface interactions, probe shape is purely a geometric factor.

When evaluating the impact of probe aperture shape on the P&P of single cells, the results show a reversed trend compared to spheroid handling. Specifically, as seen in Figure 4 D), the PBS-containing tube-tipped probes achieved a significantly lower success rate for placing single cells, approximately 50%, whereas the PBS-containing flat-tipped probes successfully placed slightly over 80% of cells. We attribute this to the tube-tipped probe’s aperture diameter, which ranges from 10 to 14 *μ*m—comparable to or larger than the diameter of the cell soma (see Figure S2). This similarity in size can hinder effective manipulation during placement, potentially causing issues such as clogging of the tube. It was noted during the experiment, that cells tended to get stuck inside the aperture and were not released again until several P&P cycles later, also compromising these attempts. The smaller (8 *μ*m) aperture diameter of the flat-tipped probe helps mitigate this challenge (see example of successful cell placement in Figure S4). Consequently, as shown in Figure 4 E), the placement rate of single cells is also significantly lower when using tube-tipped probes. These findings underscore the importance of probe aperture shape in optimising the P&P process for neurons, highlighting that this geometric aspect is crucial for achieving efficient and reliable cell placement.

### 3.2 Automating pick & place with macros for time optimisation

Neurons and their agglomerates can vary widely in size and shape, presenting challenges during P&P. These variations along with the previously-described experimental conditions can result in sub-optimal conditions, significantly increasing the time required for the process. To streamline the workflow and reduce repetitive experimental tasks, we implemented automation through macros. These macros automate several key steps: approaching the spheroid, applying negative pressure for picking, transferring the spheroid, approaching the placing surface, applying positive pressure for placing, and washing the probe every few cycles. We evaluated the efficiency of P&P for spheroids and single cells using three different approaches: the standard manual approach (without macros), the fully automated approach (with macros), and a combined approach where the user intervenes if a step in the P&P process fails.

To justify the usefulness of the automated approach, we looked at the percentage of previously targeted spheroids and single cells that were actually picked up by the probe, we examined the percentage of picked spheroids and single cells that were placed at the desired location and the speed at which this was accomplished. Targeted spheroids refer to the spheroid locations that are manually defined by the user at the beginning of the process.

In Figure 5 A), a similar success in picking the targeted spheroids for manual, macros and the combined approach is shown. This suggests that the largest factor contributing to the success rate is the proper positioning of the probe during placing, which is in all three cases done manually by the user. When looking at the success rate of placing the picked spheroids, Figure 5 B) shows that, even though macros alone do not improve the percentage, the combination of macros with the user intervention significantly increases the placing success as opposed to only macros and it is comparable with only manual P&P. Furthermore, the combined approach also improves the spheroid placing rate, making the placing twice as fast, as shown in Figure 5 C).

**Fig. 5:**
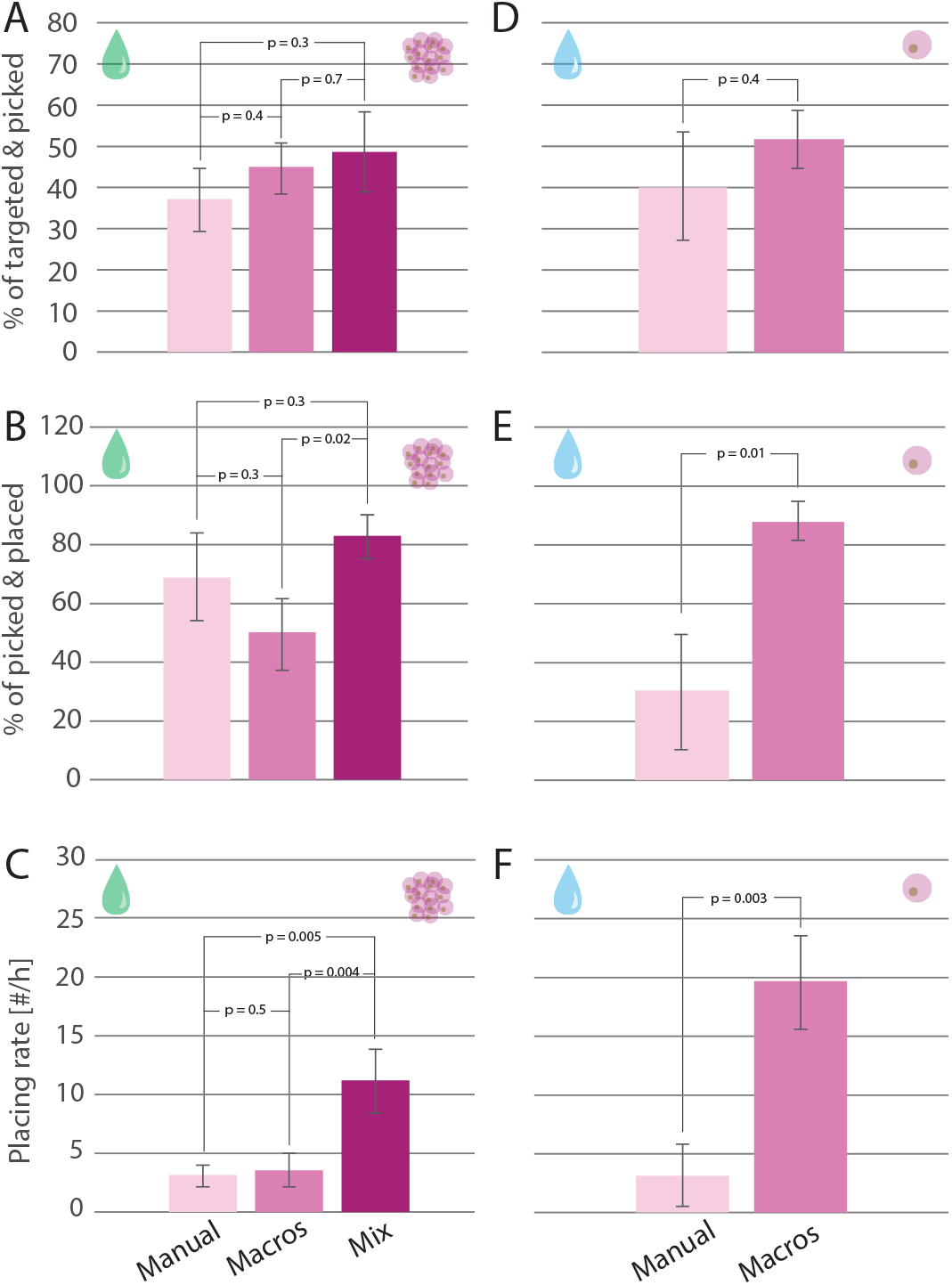
Comparison of P&P of spheroids and single cells with manual and automated execution. Percentage of targeted spheroids successfully picked (A) and placed (B) by the probe along with the rate of spheroid placement (C), the percentage of targeted single cells successfully picked by the probe (D), the percentage of picked single cells successfully placed with the probe (E), and the rate of single cell placement (F) when using manual, macros and combined approach.

In the case of single cell picking and placing, the success rate of picking the targeted cell by the probe remains consistent whether using a manual or automated approach. This consistency is attributed to the importance of precisely positioning the probe above the cell, as mentioned earlier. However, the success rate for placing cells increased nearly threefold with the use of macros, rising from approximately 30% to around 90%, as shown in Figure 5 E). Additionally, Figure 5 F) demonstrates a nearly fivefold increase in the placing rate when using macros. These results indicate that the optimal parameters for single cell P&P were effectively implemented in the macros. The significant improvement in single cell P&P performance achieved solely through automation made the combined approach, involving manual intervention, unnecessary. The entire process could be fully automated, apart from the initial selection of the target cells.

We attribute the differences in the feasibility of P&P automation between single cell and spheroid scenarios to the inherent variability in shape and size of the samples. Upon thawing or dissociation, single cells exhibit relatively uniform shape and size, making it easier to optimise handling parameters within a narrow range. In contrast, spheroids can vary significantly in both shape and size, even when users select those with similar diameters. This variation in z direction might be difficult to see and thus poses additional challenges, complicating the optimisation process for automation. Additionally. the adhesiveness of the cells to each other changes from batch to batch. While this also seems to depend somewhat on the age, it is not always consistent. Therefore, spheroids will require different parameters depending on the batch themselves.

### 3.3 Selection of cells based on fluorescence

The FluidFM OMNIUM system, used throughout the paper, is also equipped with an illumination setup suitable for fluorescence microscopy, which makes the selection of targeted spheroids based on fluorescence characteristics straightforward. In the first demonstration, we prepared spheroids containing a mixture of astrocytes and RFP-positive cortical neurons. The system allows for distinguishing between spheroids with multiple neurons and those with a single neuron by analysing fluorescence intensity and spatial localisation, as illustrated in Figure 6 A).

**Fig. 6:**
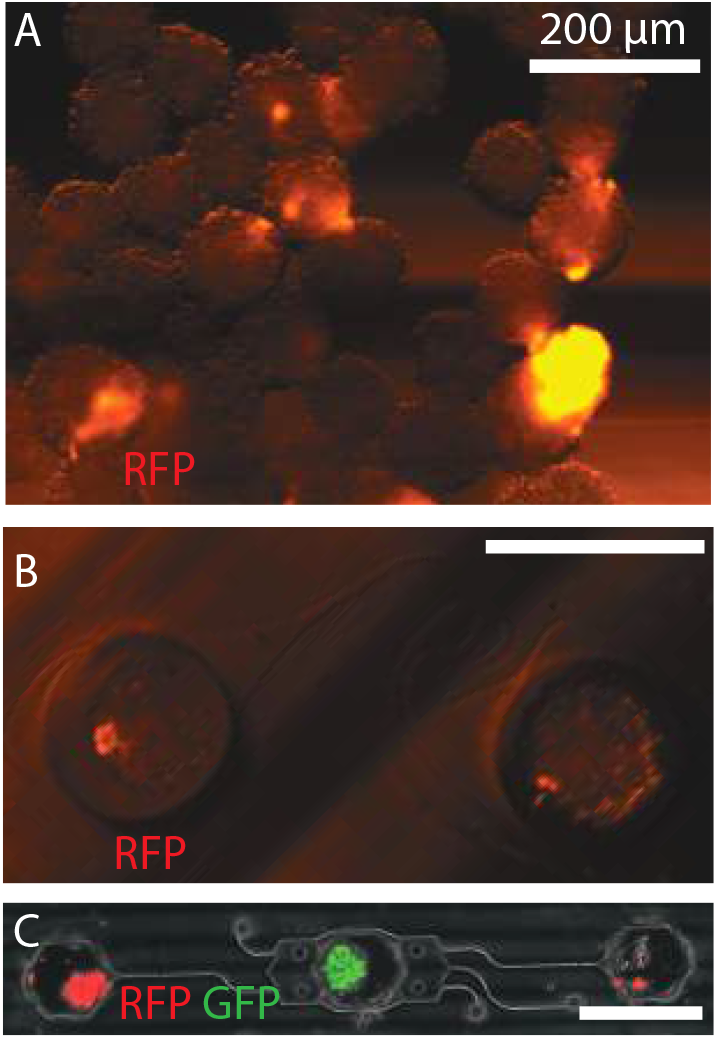
Fluorescence-based sorting. A) Spheroid mixtures of fluorescent neurons and non-fluorescent astrocyte cultures. B) Spheroids with one neuron per spheroid were picked and placed inside PDMS wells. C) Example of fluorescent-based sorting where GFP-expressing neurons were placed in the middle well and RFP-expressing neurons were placed in the side wells.

Spheroids exhibiting low fluorescence intensity concentrated in a small region, indicating the presence of only a single neuron were specifically targeted and picked. These selected spheroids were then placed into designated PDMS wells (Figure 6 B)), thereby creating a controlled network of single, interconnected neurons with supporting astrocytes. Spheroids that were placed inside the PDMS wells were imaged 12 days upon seeding and we observed healthy morphology and axon projections (see Figure S1).

Furthermore, as shown in Figure 6 C), we demonstrate the placement of a GFP-expressing neuronal spheroid into a central PDMS well and RFP-expressing spheroids into adjacent wells. In this configuration, the side wells act as ’input’ presynaptic nodes, with microchannels directing axons toward the central ’output’ postsynaptic node. This setup exemplifies how to culture spheroids from two different cell types, establishing a co-culture with directed connectivity and controlled information propagation.

### 3.4 Pick and place of neurons possible in a variety of substrates

Finally, we show the ability to successfully place neurons on different substrates. This demonstrates not only the multimodality and adaptability of the FluidFM technique, but also the potential to fine tune the P&P parameters to obtain viable and controlled networks on different surfaces for different purposes, from glass substrates to MEAs for assessing morphological and functional network characteristics.

In Figure 7 A), we show neural spheroids placed on a glass bottom surface to form a specific pattern (in this case the abbreviation of our laboratory). Similar thing has been demonstrated with single cells (See Figure S4). The placement example can be seen in the supplementary video. This example shows the level of precision and control we can achieve for the spheroid placement. The spheroids were picked and placed in a period of 85 min. In Figure 7 B) we observe a PDMS microstructure consisting of ten independent networks. Each network contains a total of three seeding wells, two being presynaptic or an ’input’ well, and one being postsynaptic or an ’output’ well. All of the 30 wells were filled with spheroids using semi-automated P&P macros and a flat-tipped probe over a period of 132 min. In Figure 7 C) we show an example of a MEA surface with PDMS microstructures being filled with spheroids using semi-automated P&P macros tool. This shows the potential of the P&P for different applications, whether the experimenter is interested in examining cell morphology and aims for creating reproducible networks on microscopy glass slides or they have interest in studying network connectivity and functional properties, therefore placing them on substrates such as MEAs.

**Fig. 7:**
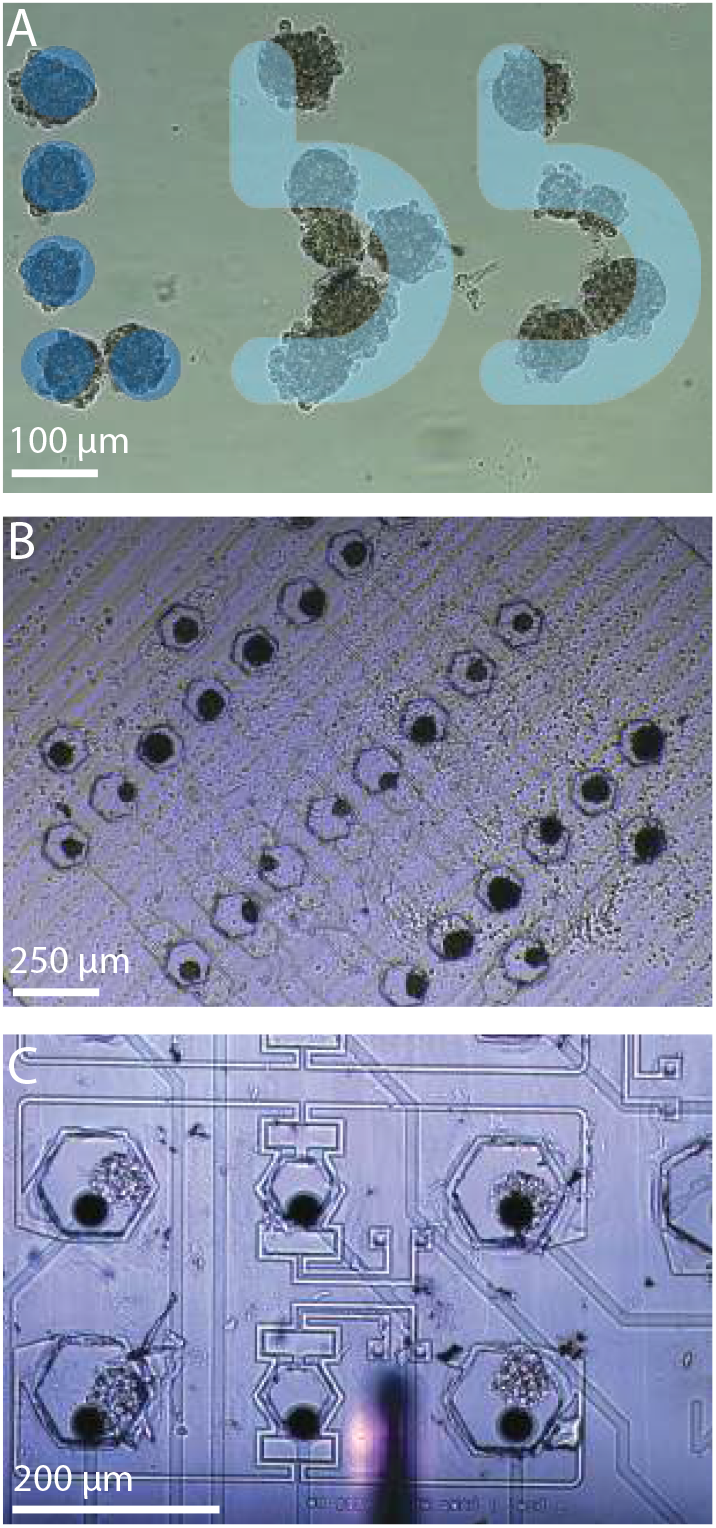
Placing neurons on a variety of substrates. A) Placing spheroids on a glass substrate in a defined pattern. B) Placing spheroids into PDMS microwells. C) Placing spheroids into PDMS microwells on a MEA. The out-of-focus FluidFM cantilever is also visible in the middle at the bottom of the image.

## 4 Conclusions

In this work, a versatile tool for both fully automatic and semi-automatic P&P of single neurons and neural spheroids, aimed at conducting reproducible cell experiments is introduced. Single cells can be placed at a rate of nearly 20 cells per hour, while spheroid placement is achieved at a rate of 11 spheroids per hour. To optimise the P&P process and expand its applicability, various probe parameters were examined to determine their impact on performance.

The findings reveal that among probe filling, probe coating, and probe shape, the shape of the probe exerts the greatest influence on performance. Furthermore, the requirements for P&P differ between single cells and spheroids. Single cell P&P is most effective with a flat-tipped probe, whereas spheroid P&P is significantly enhanced when using a tube-tipped probe.

The successful automation of the P&P process was demonstrated, allowing researchers to minimise time spent on culture preparation. For spheroids, the optimal approach is semi-automated, with user assistance at certain steps to accommodate the wider variability in spheroid sizes and shapes, which necessitates a broader range of suitable P&P parameters. In contrast, single cell P&P can be fully automated with a high success rate, thanks to the higher uniformity in single cell shape and size.

The FluidFM OMNIUM system, equipped with an incubator for long-term experiments, maintains cell viability and, when paired with macros software for (semi-)automated manipulation, allows P&P experiments to be conducted remotely from another PC or in parallel with other tasks with minimal supervision. This flexibility enhances user convenience.

Combined with fluorescent-based sorting, the P&P tool offers a powerful and flexible method for creating well-defined, reproducible *in vitro* neuronal networks, facilitating the controlled co-culture of different cell types.

There is room for improvement in the P&P system to enhance the viability of neuronal networks. It is crucial to distinguish healthy cells from dying ones, especially since iPSC-derived neurons can exhibit a high percentage of dying cells immediately after thawing. A potential solution is to develop a machine learning algorithm capable of distinguishing live from dead or dying neurons (see Figure S3) and implement it in the existing software. By selecting neurons classified as live, the overall health of the cultures can be improved.

To expedite the P&P process, optimising the selection of target positions for picking is essential. Reducing the total experiment time could also enhance the effectiveness of the macros, as it minimises the likelihood of cells shifting positions on the substrate, which can make initially selected positions sub-optimal. Additionally, exploring the optimal age for spheroid handling is worthwhile, as preliminary experiments indicate that two days post-preparation in the AggreWell may be ideal. Another area of improvement is identifying an antifouling coating for the probe that offers a higher P&P success rate and placement precision compared to non-coated probes. Moreover, consistently achieving precise single-cell placement within PDMS wells remains a challenge.

Overall, P&P using the FluidFM OMNIUM system offers several key advantages for conducting successful *in vitro* experiments. The optical selection of cells enables the choice of cell types based on fluorescent markers or desired shapes. This is particularly relevant for iPSC-derived neurons, which often show low initial viability and can differentiate into multiple undesired cell types, necessitating some form of selection or filtering. Additionally, different cell types can be selected from various locations and combined into a defined co-culture network with a precise ratio. This single-cell level control contributes significantly to the reproducibility of experiments. Moreover, the submicron placement precision of single neurons and spheroids is well-suited for the emerging field of constructing well-defined microphysiological systems.

## Author Contributions

S.C., K.V. and B.C. optimised the spheroid picking and placing protocol. K.V., J.D. and T.R. designed the PDMS microstructures for the experiments. S.C., K.V., J.D. and T.R cultured and prepared spheroids and prepared PDMS structures and dishes for experiments. E.Z.E. and S.C. inspected and prepared tube tipped-probes for picking and placing. S.C. and M.S. conducted picking and placing experiments. S.C. and K.V. conducted results analysis and writing. J.V. supervised the study, revised the manuscript and is the corresponding author. All authors read and approved the manuscript.

## Acknowledgements

The research was financed by ETH Zurich, Innosuisse (Project No. 55051.1 IP-ENG), the Swiss National Science Foundation (Project Nr: 165651 and CRSII5 202301 / 1), Eurostar (Project Nr: Eurostar E!11644 SOUL) and the Human Frontiers Science Program Organization, HFSPO. The authors would like to thank Cytosurge for their technical support. The authors also thank Novartis for generously providing hiPSC neurons. The authors further thank Edin Sarajlić (Bruker Nederland BV, Netherlands) for designing and fabricating tube-tipped probes.

