## Supplementary information for "Constructing well-defined neural networks of multiple cell types by picking and placing of neuronal spheroids using FluidFM"

#### 1 ARYA Macros for Rat Primary and hiPSC-derived Spheroid Pick and Place

1. Select Position
   - Coloured: Blue
   - Select one or more picking points
2. Select Position
   - Coloured: Yellow
   - Select one or more placing points
3. FluidFM Calibration
   - Coloured: Blue
   - Configuration:
     - To Position:
       * Target Position: (Bottom of Well + 0.25) mm
       * Speed: 500 $\mu$ms$^{-1}$
       * Pressure Mode: Keep
     - Move Distance:
       * Delta: -100 $\mu$m
       * Speed: 400 $\mu$ms$^{-1}$
       * Pressure Mode: Keep
     - To Setpoint:
       * Setpoint: 25 mV
       * Speed: 20 $\mu$ms$^{-1}$
       * Pressure Mode: Keep
     - Keep Setpoint:
       * Setpoint: 25 mV
       * Duration: 2 s
       * P: -0.001
       * I: -0.0005
       * D: 0
       * Pressure Mode: Set
       * Pressure: -500 mbar
     - Move Distance:
       * Delta: 10 $\mu$m
       * Speed: 20 $\mu$ms$^{-1}$
       * Pressure Mode: Keep
     - Keep Position:
       * Duration: 1 s

        * Pressure Mode: Set
        * Pressure: -50 mbar
  - intra well xy speed: 4 mms$^{-1}$
  - Record Curve: No

4. FluidFM Calibration
  - Coloured: Blue
  - Configuration:
    - Move Distance:
      * Delta: 20 $\mu$m
      * Speed: 40 $\mu$ms$^{-1}$
      * Pressure Mode: Keep
    - Keep Position:
      * Duration: 1 s
      * Pressure Mode: Keep
    - Move Distance:
      * Delta: 100 $\mu$m
      * Speed: 200 $\mu$ms$^{-1}$
      * Pressure Mode: Keep
    - Keep Position:
      * Duration: 1 s
      * Pressure Mode: Keep
    - Move Distance:
      * Delta: 400 $\mu$m
      * Speed: 800 $\mu$ms$^{-1}$
      * Pressure Mode: Keep
  - intra well xy speed: 4 mms$^{-1}$
  - Record Curve: No

5. FluidFM Calibration
  - Coloured: Yellow
  - Configuration:
    - To Position:
      * Target Position: (Bottom of Well + 0.25) mm
      * Speed: 400 $\mu$ms$^{-1}$
      * Pressure Mode: Keep
    - To Setpoint:
      * Setpoint: 50 mV
      * Speed: 80 $\mu$ms$^{-1}$
      * Pressure Mode: Set
      * Pressure: 200 mbar
    - Keep Position:
      * Duration: 20 s
      * Pressure Mode: Set
      * Pressure: 500 mbar
    - Move Distance:
      * Delta: 20 $\mu$m
      * Speed: 40 $\mu$ms$^{-1}$
      * Pressure Mode: Keep
  - intra well xy speed: 4 mms$^{-1}$
  - Record Curve: No

6. Move XYZ 1.07.42
   - Coloured: Yellow
     - XY movement: Selected point
     - X offset from point: 0 $\mu$m
     - Y offset from point: -30 $\mu$m
     - Z movement: Content height
     - Z offset from content height: 0 $\mu$m
     - Focus option: Relative to current focus
7. FluidFM Calibration
   - Coloured: Yellow
   - Configuration:
     - Move Distance:
       * Delta: 100 $\mu$m
       * Speed: 4 $\mu$ms$^{-1}$
       * Pressure Mode: Set
       * Pressure: 200 mbar
     - Move Distance:
       * Delta: 400 $\mu$m
       * Speed: 4 $\mu$ms$^{-1}$
       * Pressure Mode: Keep
   - intra well xy speed: 4 mms$^{-1}$
   - Record Curve: No
8. FluidFM Calibration
   - Coloured: Blue
   - Configuration:
     - To Position:
       * Target Position: (Bottom of Well + 0.25) mm
       * Speed: 4 mms$^{-1}$
       * Pressure Mode: Keep
   - intra well xy speed: 12 mms$^{-1}$
   - Record Curve: No

#### 2 ARYA Macros for Single hiPSC-derived Neuron Pick and Place

1. Select Position
   - Colour: Yellow
   - Select picking points
2. Select Position
   - Colour: Blue
   - Select placing points
3. FluidFM Calibration
   - Colour: Yellow
   - Configuration:
     - To Position
       * Target Position: (Bottom of Well + 0.1) mm
       * Speed: 16 mm/s
       * Pressure Mode: Set
       * Pressure: 1 mbar
     - intra-well xy speed 32 mm/s
     - Record Curve: No
4. Correct Position
   - Colour: Yellow
   - Z position option: ContentHeightOrLower
   - X offset of tip during detection: 0 $\mu$m
   - Y offset of tip during detection: -30 $\mu$m
   - Immersion Speed: 10 mm/s
   - Settings to apply during idle time: Current settings
5. FluidFM Calibration
   - Colour: Yellow
   - Configuration:
     - To Setpoint:
       * Setpoint: 50 nN
       * Speed: 160 $\mu$ms$^{-1}$
       * Pressure Mode: Set
       * Pressure: -50 mbar
     - Keep Position:
       * Duration: 1 s
       * Pressure mode: Set
       * Pressure: -10 mbar
     - Move Distance:
       * Delta: 10 $\mu$m
       * Speed: 160 $\mu$ms$^{-1}$
       * Pressure Mode: Keep
     - Move Distance:
       * Delta: 20 $\mu$m
       * Speed: 320 $\mu$ms$^{-1}$
       * Pressure Mode: Keep
     - Move Distance:
       * Delta: 100 $\mu$m
       * Speed: 1.6 mms$^{-1}$

    * Pressure Mode: Keep
  - Keep Position:
    * Duration: 1 s
    * Pressure Mode: Keep
- intra well xy speed: 32 $mms^{-1}$
- Record Curve: No

6. FluidFM Calibration
   - Colour: Yellow
   - Configuration:
     - Move Distance:
       * Delta: 1.5 mm
       * Speed: 6.4 $mms^{-1}$
       * Pressure Mode: Keep
   - intra-well xy speed: 32 $mms^{-1}$
   - Record Curve: No

7. FluidFM Calibration
   - Colour: Blue
   - Configuration:
     - To Position:
       * Target Position: (Bottom of Well + 0.1) mm
       * Speed: 3.2 $mms^{-1}$
       * Pressure Mode: Keep
     - To Setpoint:
       * Setpoint: 50 nN
       * Speed: 160 $\mu ms^{-1}$
       * Pressure Mode: Keep
     - Keep Position:
       * Duration: 5 s
       * Pressure Mode: Set
       * Pressure: 30 mbar
     - Move Distance
       * Delta: 20 $\mu$m
       * Speed: 320 $\mu ms^{-1}$
       * Pressure Mode: Keep
     - Move Distance:
       * Delta: 100 $\mu$m
       * Speed: 32 $mms^{-1}$
       * Pressure Mode: Keep
     - Move Distance
       * Delta: 1.2 mm
       * Speed: 32 $mms^{-1}$
   - intra-well xy speed: 32 $mms^{-1}$
   - Record Curve: No

8. Wash
   - Current count: 0
   - Interval (0 = never): 3
   - Return to current position after washing: Yes
     - B3: 1000 mbar, 30 s [Tergazyme]

- B2: 1000 mbar, 5 s [PBS]
- C2: 1000 mbar, 5 s [PBS]
- A4: 1000 mbar, 5 s [Ultrapure water]
- B4: 1000 mbar, 5 s [Ultrapure water]
- C4: 200 mbar, 5 s [Ultrapure water]

#### 3 Supplementary Figures

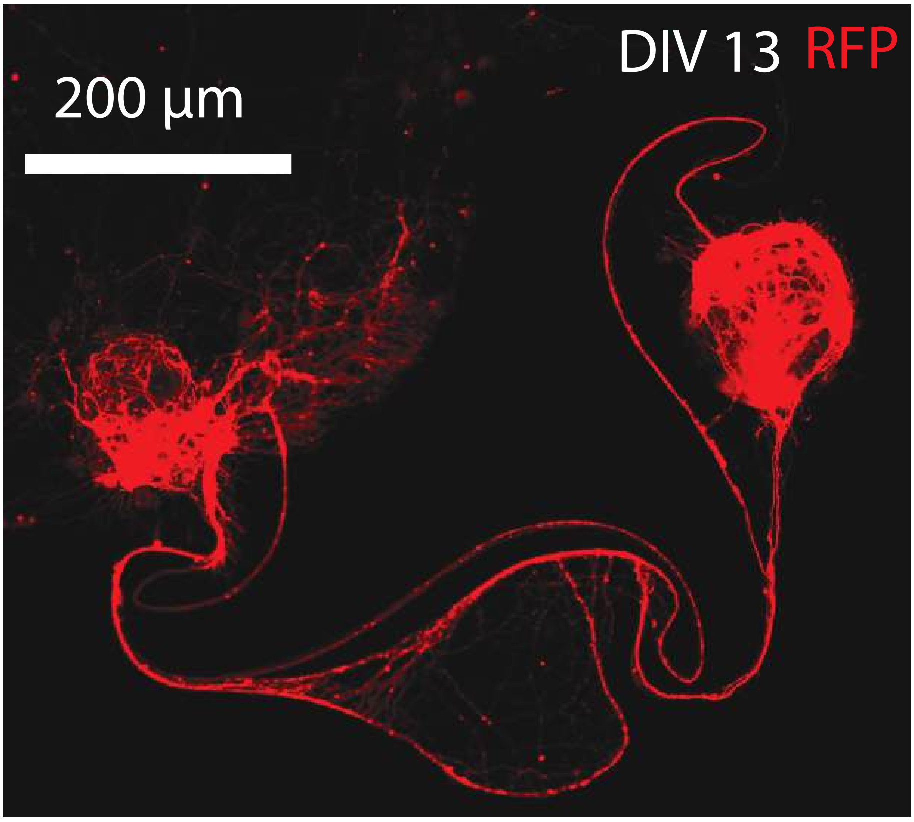


Figure S1: Spheroids placed within PDMS wells using the FluidFM OMNIUM system were imaged at DIV13. These spheroids contain a mix of RFP-expressing NGN2 neurons and astrocytes.

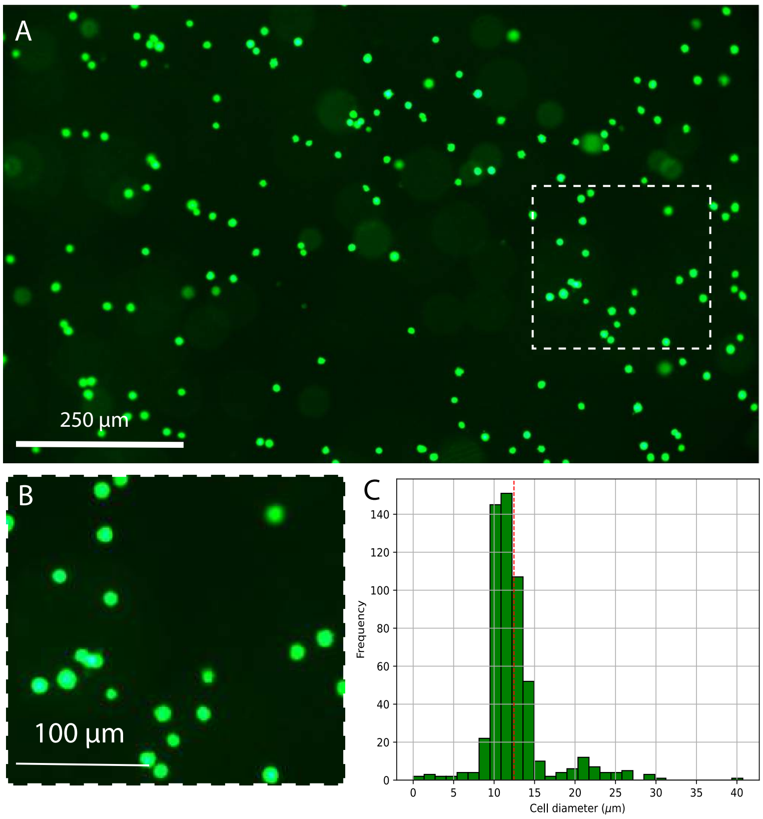


Figure S2: A) hiPSC-derived neurons suspended on the picking surface directly upon thawing. B) Zoom in. C) Distribution of hiPSC-derived neuron diameter directly upon thawing.

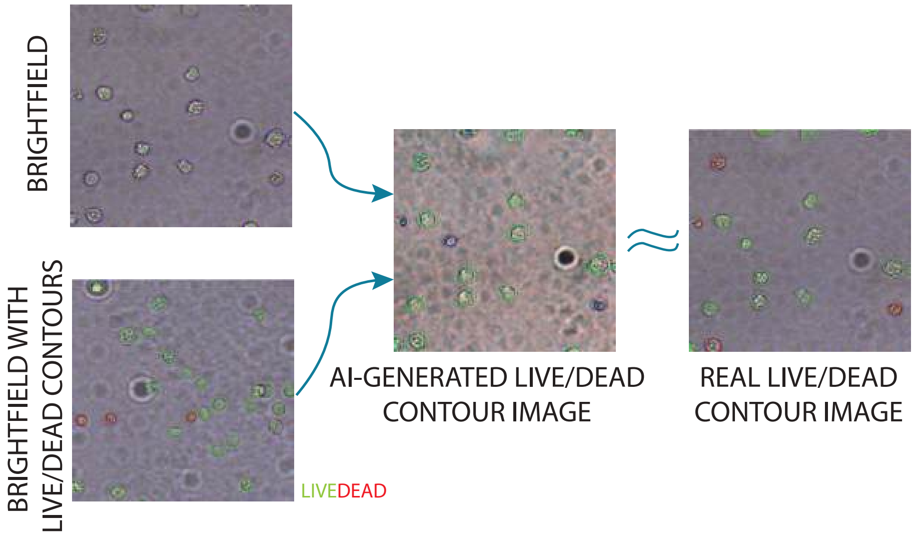


Figure S3: Idea of automated selection of live cells for picking using an ML tool.

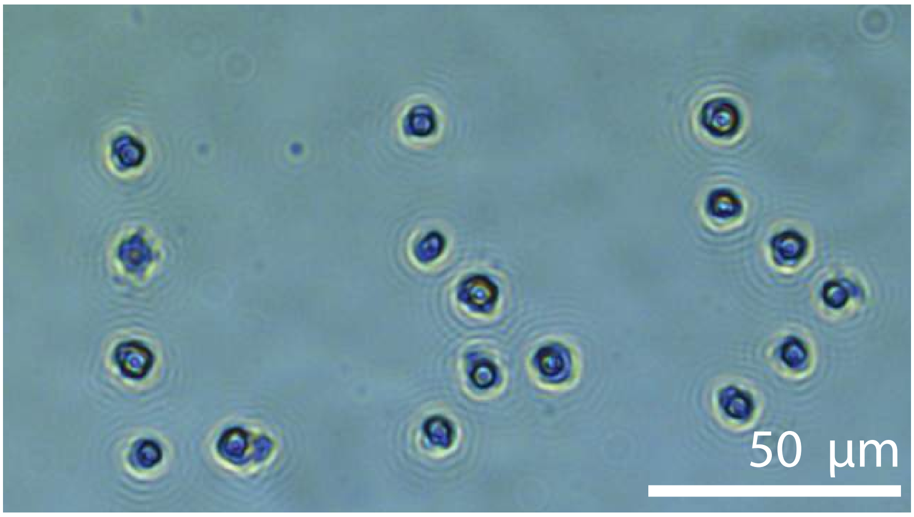


Figure S4: Example of single cell placement. The P&P was performed using a flat, PBS-filled probe.

### 4 Supplementary Videos

Supplementary videos may be found here.
